# Machine learning approaches for classification of Plasmodium falciparum life cycle stages using single-cell transcriptomes

**DOI:** 10.1101/2022.06.22.497155

**Authors:** Swarnim Shukla, Soham Choudhuri, Gayathri Priya Iragavarapu, Bhaswar Ghosh

## Abstract

Malaria, spread by the female *Anopheles* mosquito, is a highly fatal disease widespread in many parts of the world, causing 0.4 million deaths globally. Vital gene expressions form the basis in the detection of malaria infection levels. Quantification of malaria parasite infected RBCs and classification of its life cycle stages are done at macroscopic level by experts, for making informed decisions. Off late multiple computational approaches have been proposed to circumvent the problem of dimensionality leading to accurate predicted results. In this work a dimensionality reduction technique based on Genetic Algorithm (GA) is applied on *P. falciparum* single-cell transcriptomics to arrive at an optimized subset of features from the larger dataset. Features are chosen based on their class variants considering increased efficiency and accuracy, to separately transform the selected elements into a lower dimension. For the classification of the life cycle of malaria parasite based on single cell transcriptome data, a three-pronged approach employing the multiclass Support Vector Machine (SVM), Logistic Regression (LR) and Random Forest (RF) techniques is used. Distribution of cells was visualised and mapped using the R-based Seurat package. Further, we constructed protein interaction networks of the genes identified by the feature selection method and elucidated the role of the proteins in progression of the parasite through it’s life cycle. Our approach presents a novel protocol to implement ML techniques on scRNA seq datasets and subsequently harnessing the extracted information for biomarker/drug target detection.

## Introduction

Malaria is a deadly disease caused by the *Plasmodium* parasite and is transmitted through the bite of a female Anopheles mosquito. This *Plasmodium* attacks the red blood cells (RBCs) and the degree of malaria can be estimated by the quantity of infected RBCs.^1^ The complex life cycle of malaria parasites features diverse developmental strategies, each of which is uniquely adapted to thrive in the particular host environment. Six *Plasmodium* species cause human malaria, with majority of the estimated 0.4 million annual deaths caused by *Plasmodium falciparum* (*P. falciparum*). Blood stage development begins when a newly released, extracellular parasite (a merozoite) invades an erythrocyte, establishing the ring stage of infection, progressing to the trophozoite stage. During this stage the infected erythrocyte is extensively modified to enable parasite proliferation. Thereafter the parasite divides to form a connected group of daughter cells, called schizont, which eventually lyses the host erythrocyte, releasing the newly formed merozoites to invade new erythrocytes. These steps are collectively known as the intraerythrocytic developmental cycle (IDC). ^2^ Malaria symptoms include high fever and headache and in some severe cases even seizures and death are caused. Malaria mostly affects the economically weak communities of the world, where medical treatment is not readily available. For a quick and successful recovery of any patient, it is vital to diagnose and treat the malarial infection early. If the malaria life cycle stages are somehow ascertained then the treatment of disease becomes easier. Experienced medical professionals frequently examine a large number of blood films to detect malaria infection. Microscopists normally visualize the thick and thin blood smears to identify a disease or its cause. However, the accuracy depends upon smear quality and the expertise in classifying and counting the parasite and non-parasite cells. It is fairly challenging to number the parasites and infected RBCs manually and needs an expert microscopist for quality diagnosis.^3^ Recent advances in single cell RNA-sequencing (sc-RNA)techniques paved new ways to characterize gene expression changes during the development stages of the plasmodium life cycle.^4,5^ Analysis of the gene expression regulation may allow us to identity new diagnostics marker as well as potential target for a new drug. Indeed, many studies have already been conducted in the last few years by utilizing sc-RNA experiments for *Plasmodium falciparum*. One of the central advantages of employing sc-RNA methods is the scope of exploring cell to cell heterogeneity in the population by uncovering hidden variability in gene expression among single cells ^5–7^. In fact, recent studies have elucidated the role of heterogeneity in enabling a small fraction of *Plasmodium* population inside the human host to remain ready to enter into the mosquito host by making transition to gametogenesis stage.^8^ Similarly, heterogeneity plays crucial role in *Plasmodium* stress response inside the RBC.^9^ However, the diagnostics of scRNA-Seq is challenging as its outcome suffers from lack of fit due to high dimensional gene expression data. Advanced computational skills are needed to study and process the massive volume of proteomic and genomic data obtainable freely from several repositories ^10,11^ and harness them to reveal new biological insights.

The dataset contains redundant characteristics that behave as noise during model training. As a result, classification performance is degraded and computing time is increased. Dimensionality Reduction (DR) techniques are required to eliminate redundancy and to retrieve irrelevant details that hinder performance. There are two methods for reducing the dimensionality of data: Feature Extraction and Feature Selection. Feature selection is further divided into filter, wrapper, and embedded methods. ^12^ In the filter method, mathematical measures are used to select the optimal features. Wrapper is a feedback method that uses a machine-learning algorithm to help choose the best features. The embedded approach is a hybrid of the filter and wrapper methods. This paper proposes a wrapper-based feature selection technique using the Genetic Algorithm to select the optimal features and remove the redundant noise in the dataset. In order to evaluate the performance of these features, SVM, LR, and RF classification models are used.

The main objective of our study is to select top-ranked genes from the scRNA-seq profiles at different stages of the plasmodium falciparum life cycle inside infected RBC. We employ supervised learning algorithm coupled with feature selection algorithms to extract most relevant genes in predicting the life cycle stages of plasmodium inside RBC. The first stage of the proposed model is to optimize the quality data from the data set by removing the redundant, noisy, and irrelevant genes (features). From the literature review(see Discussion and outlook) it can be concluded that genetic algorithm (GA) showed that better performance than the other selection algorithms and thus, prominently used for feature selection from high dimensional data sets.

Hence, GA is employed in our proposed approach as the search algorithm in the feature selection process. This subset can be further utilized in the second stage of the process, Classification, to produce high classification accuracy. We tested the subsets using three classifiers: SVM, LR, and RF to ensure the investigation is carried out rigorously. The combination of first and second stages of the proposed model will achieve a better identification of the different Malaria Life Cycle stages. Additionally, the feature selection method is able to identify genes which significantly change expression across the life cycle stages. UMAP projection of the cells based on these features supports the distinction of stages using these features. We constructed the protein interaction network of these genes and performed a set of topological analysis to provide hierarchies according to importance of the genes in the network. These genes can be used for diagnosis and drug targets. Our study presents a theoretical framework to select diagnosis markers and drug targets by implementing ML techniques on sc-RNA-seq data.

## Results

The single cell RNA-seq dataset utilized here is derived from the Malarial Cell Atlas, an open source database of single cell transcriptomic data spanning the complete life cycle of malarial parasites. It is freely accessible through a dynamic, user-friendly web interface (www.sanger.ac.uk/science/tools/mca/mca/).^13^ For the current study, we considered the 10X scRNA-seq data of the intraerythrocytic stages of P.falciparum in the human host. The dataset has 5066 rows and 6737 columns. Each row corresponds to scRNA-seq read counts of a gene and each column corresponds the same for a single cell. There are 5066 features in this dataset, which correspond to all the genes in each cell of the parasite. Additionally, each cell is assigned one label among the four blood cycle stages (ring, early trophozoite, late trophozoite, and schizont). Thus, we set out to utilize classification ML algorithms (Material and methods) which would allow us to predict the life cycle stage of a cell based on the gene expression pattern.

### Data visualisation and dimensionality reduction of the scRNA-seq data

We used Seurat,^14^ an R-based Bioconductor package, to visualise and then apply dimensionality reduction on the single cell RNA-seq data. We integrated the raw expression counts and metadata generated by Howick V.M. et al. ^13^, for downstream analysis to visualize the cells on a suitable manifold. Since the published data had already undergone quality control, the cell counts were subjected to normalisation using the ‘LogNormalize’ method of the Seurat package. This involves a global normalisation of cell counts with respect to the total expression, followed by log transformation. For further analysis, it is useful to focus on genes that exhibit high variation over all cells in the dataset. Hence,we selected 1000 highly variable features (genes) from the data using the FindVariableFeatures() function. The data was then subjected to scaling before applying standard dimensionional reduction techniques like PCA and UMAP. Next, PCA was performed on the data and the clusters produced from this linear dimensional reduction were annotated based on the blood cycle stages. Using the first 10 PCs, we also performed a non-linear UMAP-based dimensional reduction on the cells for a better projection and annotated the clusters based on blood cycle stage. RunUMAP() function was used with dims=1:10. Figure 1 represents the UMAP projection and we notice four clusters for ring, early trophozoite, late trophozoite and schizont respectively.

**Figure 1:**
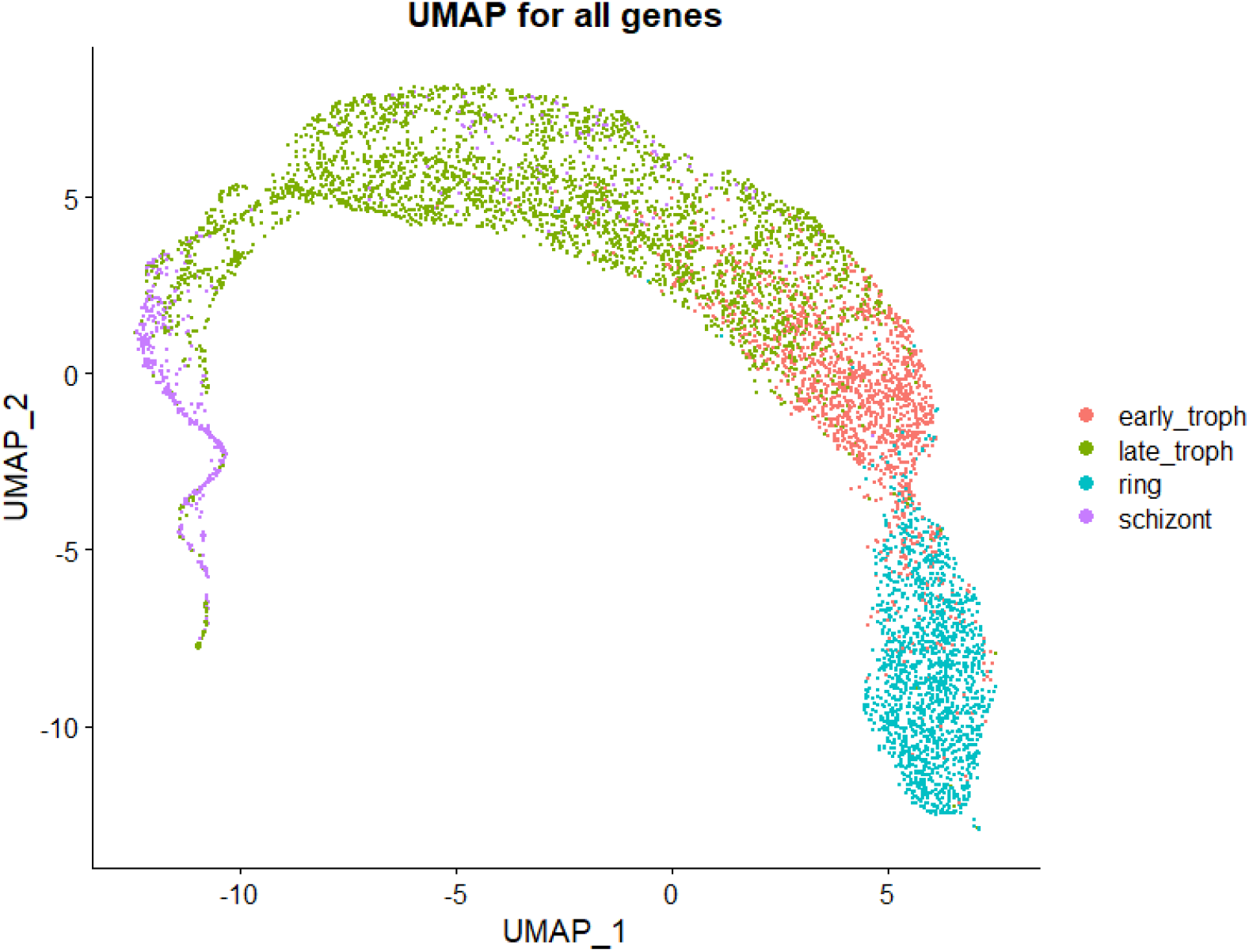
Two dimensional visualization of the scRNA data set shows distinct cluster of life cycle stages. UMAP of scRNA-seq counts of all 5066 features. Non-linear dimensional reduction of expression values of all genes in the dataset.The cell clusters are colored based on the blood cycle stages of *P*.*falciparum*.

### Classification without feature selection

Next, we implement SVM, LR and RF algorithms to classify the cells into the four different stages by including all the genes. In order to calculate the prediction accuracy, we randomly selected 80 % of the cells as training set and 20 % of the cells are chosen for calculating the desired accuracy. Figure 2 shows the accuracy of SVM, LR, and RF. This is the baseline for our experiment. Without Feature selection, SVM performed best with classification accuracy measured this 91%. The least accurate algorithm was Random Forest (88%) whilst the level of performance of LR (90%) was acceptable.

**Figure 2:**
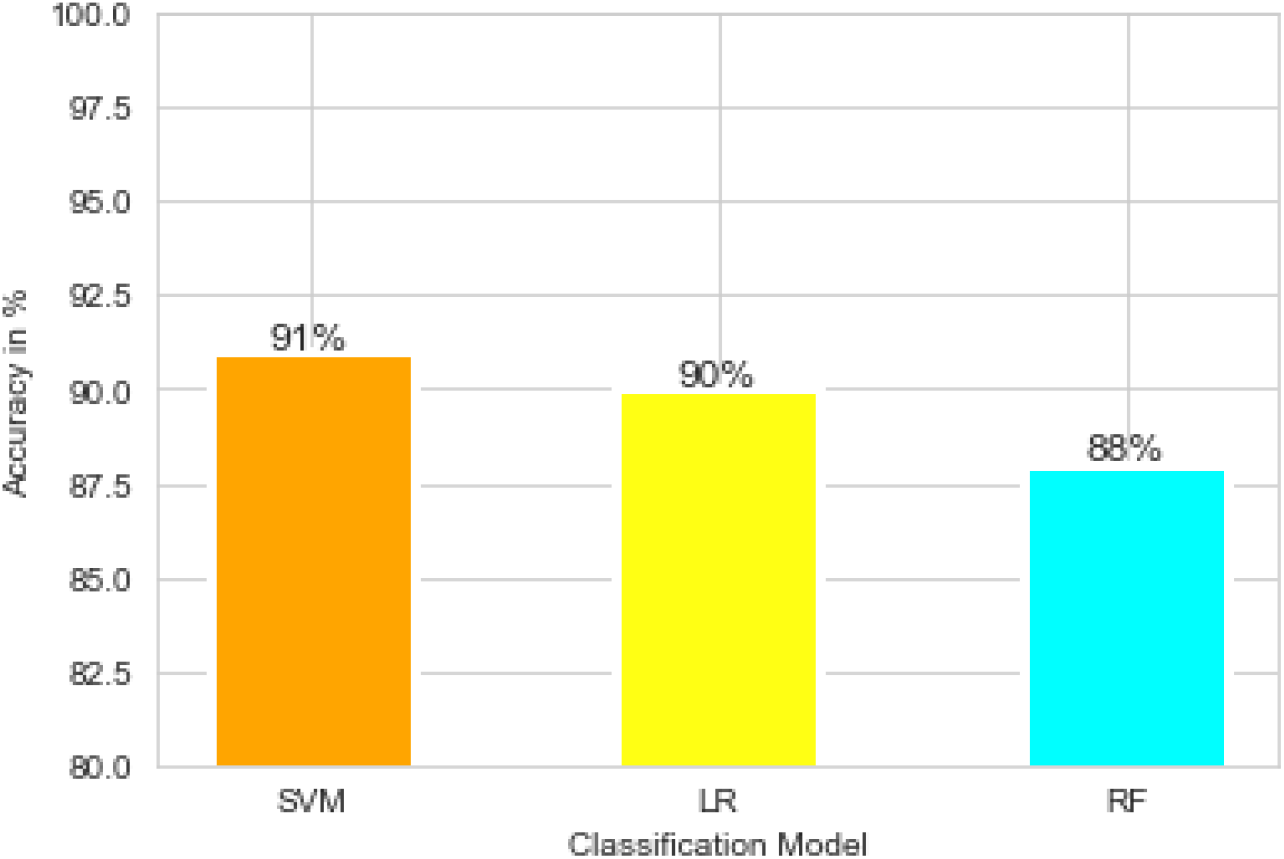
Classification accuracy of different models without feature selection. The classification accuracy are shown for different machine learning protocols namely SVM, LR and RF

As listed in Table 1, the following best F1 scores were determined: ring 96%, late troph 94%, early troph 86% and schizont 80%.

**Table 1:**
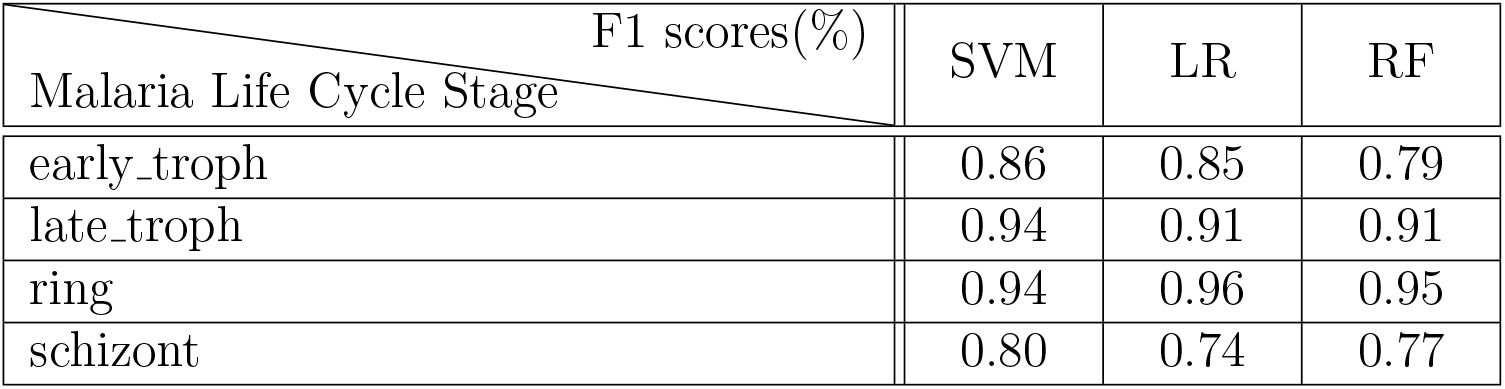
F1 scores of different models of the different classes without Feature Selection

### Features selection

From the data set, we observed that there are many genes which do not change expression levels across the life cycle stages. Thus, a feature selection algorithms would drastically reduce the dimension and would allow us to extract important genes, the expression of which are responsible for the life cycle changes. In order to select the features, a genetic algorithm (GA) based– pipeline is implemented (Material and methods for details). The GA pipeline outputs the most optimal features and has removed the redundant features. Table 2 shows the details of the number of features. The initial number of features was **5066** out of which a subset of size **336** was selected using the GA. The dataset was reduced by **93.3%**.

**Table 2:** Numbers of Features selected

|  | Number of Features |
| --- | --- |
| Full Dataset | 5066 |
| Features Selected after GA pipeline | 336 |

### Classification with feature selection

This section shows the detail of the classification results with feature selection stage. Figure 3 shows the accuracy of SVM, LR, and RF. SVM performed best with classification accuracy measured this 91%. RF and LR both gave this 90% accuracy. The coming sections shows the test results of the SVM, LR, and RF models respectively.

**Figure 3:**
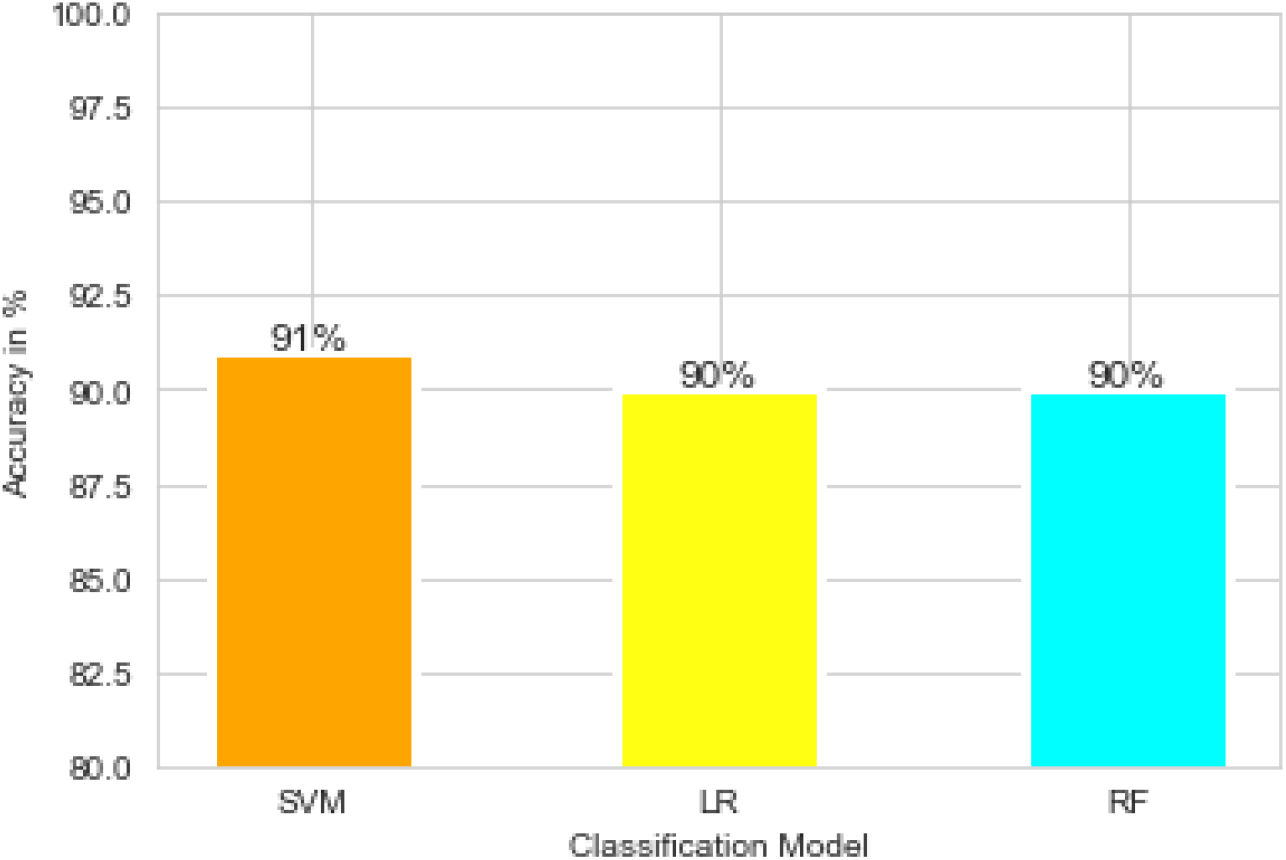
Classification accuracy of different models with feature selection. The classification accuracy are shown for different machine learning protocols namely SVM, LR and RF after selection of the 336 feature following genetic algorithm.

### Using multiclass Support Vector Machine

Table 3 presents the precision, recall, and f1 scores of the SVM model for the four different classes. For late_troph and ring, we have achieved an f1 score of 0.93 and 0.94, respectively. We got an average f1 score for early_troph at 0.86. Schizont was the worst, at 0.78.

**Table 3:** Test results of SVM model with feature selection

| Malaria life cycle stage | Metric (%) |  |  |
| --- | --- | --- | --- |
|  | precision | recall | F1-score |
| early_troph | 0.92 | 0.82 | 0.86 |
| late_troph | 0.91 | 0.95 | 0.93 |
| ring | 0.90 | 0.99 | 0.94 |
| schizont | 0.92 | 0.67 | 0.78 |

### Using Logistic Regression

Table 4 presents the precision, recall, and f1 scores of the LR model for the four different classes. For late_troph and ring, we have achieved an f1 score of 0.91 and 0.95, respectively. We got an average f1 score for early_troph at 0.87. Schizont was the worst, at 0.74.

**Table 4:** Test results of LR model with feature Selection

| Malaria Life Cycle Stage \ Metric (%) | precision | recall | F1-score |
| --- | --- | --- | --- |
| early_troph | 0.87 | 0.86 | 0.87 |
| late_troph | 0.90 | 0.93 | 0.91 |
| ring | 0.95 | 0.96 | 0.95 |
| schizont | 0.83 | 0.66 | 0.74 |

### Using Random Forest

Table 5 presents the precision, recall, and f1 scores of the RF model for the four different classes. For late_troph and ring, we have achieved an f1 score of 0.92 and 0.95, respectively. We got an average f1 score for early_troph at 0.83. Schizont was the worst at, 0.75.

**Table 5:** Test results of RF model with feature selection

| Malaria life cycle stage \ Metric (%) | precision | recall | F1-score |
| --- | --- | --- | --- |
| early_troph | 0.91 | 0.77 | 0.83 |
| late_troph | 0.88 | 0.97 | 0.92 |
| ring | 0.93 | 0.97 | 0.95 |
| schizont | 0.93 | 0.63 | 0.75 |

### Confusion matrix and mutual information between predicted and true labels for three models

Figure 4 shows the confusion matrix of the three models. The confusion matrix for the SVM model shows that 32 samples were predicted as late_troph which should have been labelled as early_troph. Similarly, 37 samples were predicted as late_troph which were otherwise labelled as schizont. For late_troph class, 15 samples were misclassified as early_troph. For ring class, 2 samples were misclassified as early_troph.

**Figure 4:**
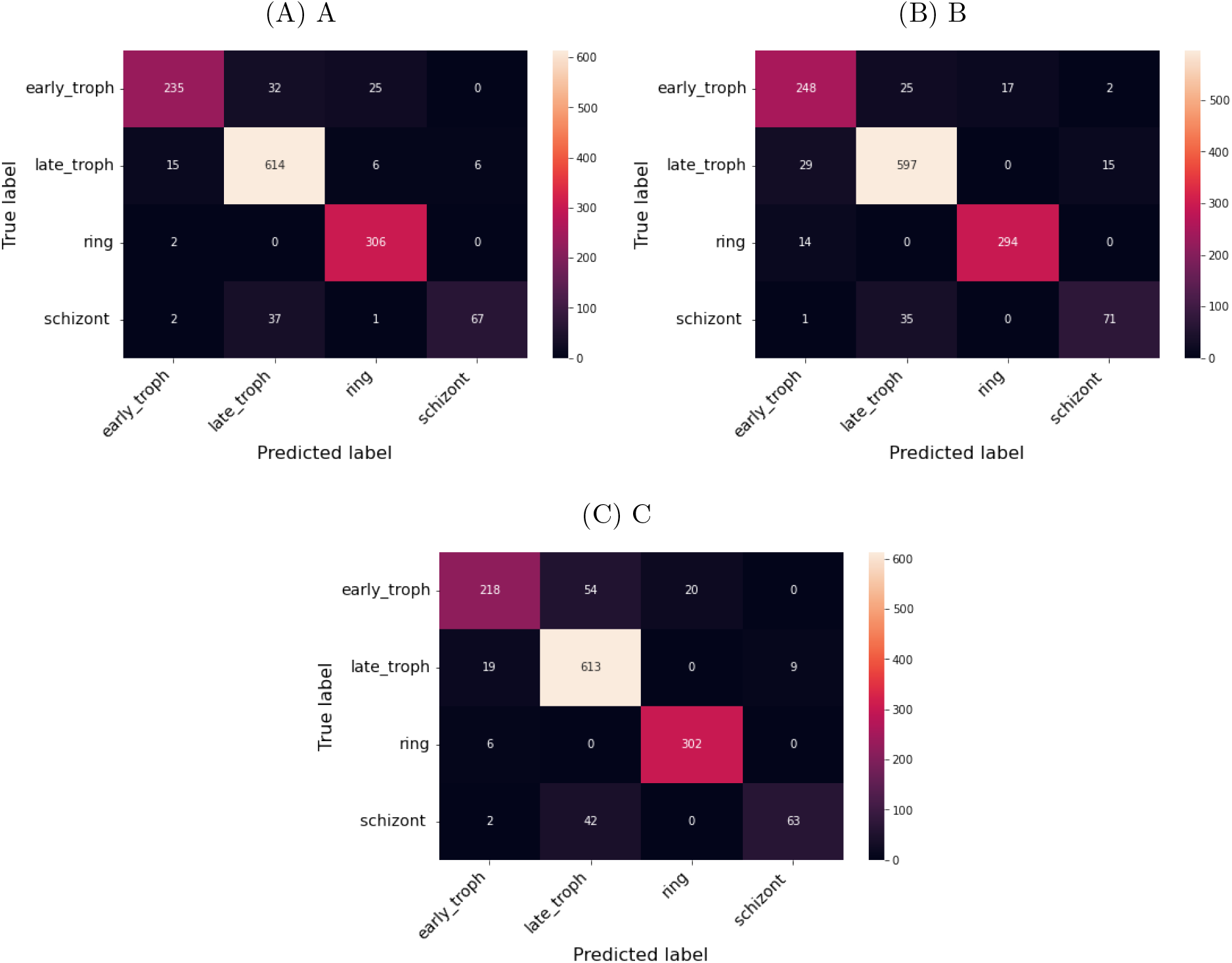
Confusion matrix of different models show the prediction accuracy for different stages. The heatmaps display the confusion matrix in predicting the four different stages as indicated after feature selection for three different models (A) SVM (B) LR (C) RF models

The confusion matrix for the LR model shows that 25 samples were predicted as late_troph which should have been labelled as early_troph. Similarly, 35 samples were predicted as late_troph which were otherwise labelled as schizont. For late_troph class, 29 samples were misclassified as early_troph. For ring class, 14 samples were misclassified as early_troph.

The confusion matrix for the RF model shows that 54 samples were predicted as late_troph which should have been labelled as early_troph. Similarly, 42 samples were predicted as late_troph which were otherwise labelled as schizont. For late_troph class, 19 samples were misclassified as early_troph. For ring class, 6 samples were misclassified as early_troph. These were some of the common misclassifications in all three models.

### A comparison of classification without vs with feature selection

Figure 5 shows the accuracy of classification without feature selection vs. classification with feature selection. We can see that we have reduced our feature set from 5066 to 336. Using these 336 features, we have achieved an accuracy of 91% in the SVM model, 90% in the LR model, and 90% in the RF model. We have improved the accuracy of the RF model and have achieved similar results in the SVM and LR models.

**Figure 5:**
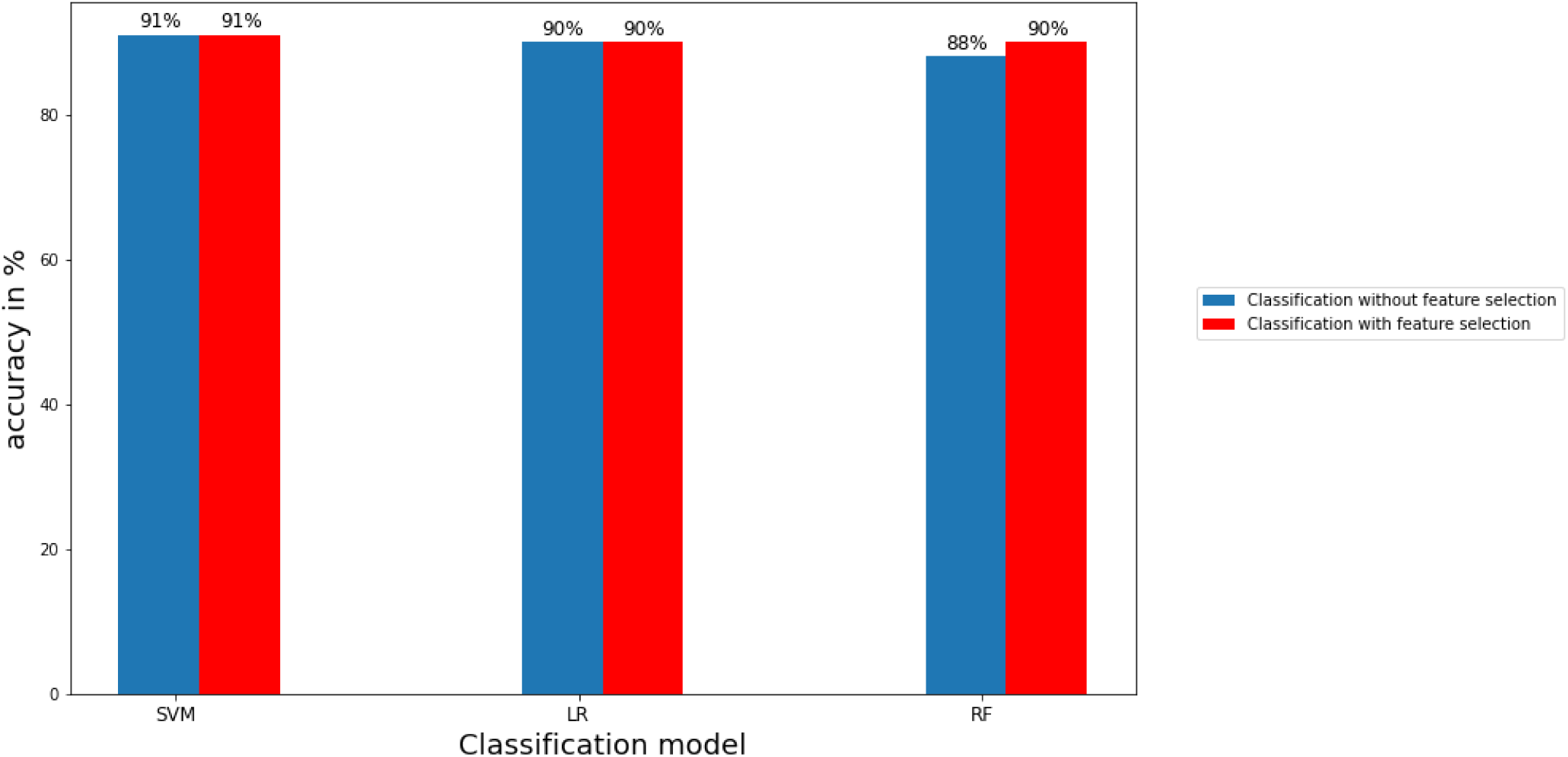
Classification accuracy with feature selection vs without feature selection demonstrate the legitimacy of the selected features. The bar graphs display a comparison between the values of accuracy for the three models and for classification with feature selection and without feature selection as indicated

For the SVM model, without feature selection, we got a f1 score of 0.86, 0.94, 0.94, and 0.80, where as with feature selection, we got a f1 score of 0.86, 0.93, 0.94, and 0.78 for early_troph, late_troph, ring, and schizont, respectively. For the LR model, without feature selection, we got a f1 score of 0.85, 0.91, 0.96, and 0.74, where as with feature selection, we got a f1 score of 0.87, 0.91, 0.95, and 0.74 for early_troph, late_troph, ring, and schizont, respectively. For the RF model, without feature selection, we got a f1 score of 0.79, 0.91, 0.95, and 0.77, where as with feature selection, we got a f1 score of 0.83, 0.92, 0.95, and 0.75 for early_troph, late_troph, ring, and schizont, respectively. We can see from the results that just by using the 336 features we have achieved similar f1 scores for all the four classes across all three models. This proves the robustness of the features selected from the GA pipeline. For the early_troph class, we have achieved the best f1 score of 0.87 from the LR model. For the late_troph class, we have achieved the best f1 score of 0.93 from the SVM model. For the ring class, we have achieved the best f1 score of 0.95 from both the SVM and RF models. Schizont was the worst, with an f1 score of 0.78 from the SVM model. This could be because less number of instances of the schizont class in the dataset compared to other classes.

We also calculated the mutual information (MI) between the predicted labels and the true labels of the three models using the joint probabilities from confusion matrix (see Subsection). For instance, C(1,1) of the confusion matrix represents the the joint probability P(X,Y) where X= true label of early_troph and Y corresponds to correctly predicted early_troph. Similarly C(1,2) would reflect the joint probability P(X,Y) where X= true label of early_troph while Y= incorrectly predicted to be late_troph. Figure 6 shows the comparison of MI with and without feature selection. One of the advantages of displaying the accuracy using mutual information is that the upper limit of the mutual information is exactly known. So, the accuracy of the model can be compared with the ideal case. In our case, since the number of labels is four, the maximum possible mutual information for an error free case is 2 bits, however maximum information acquired by the models is 1.28 bits here.

**Figure 6:**
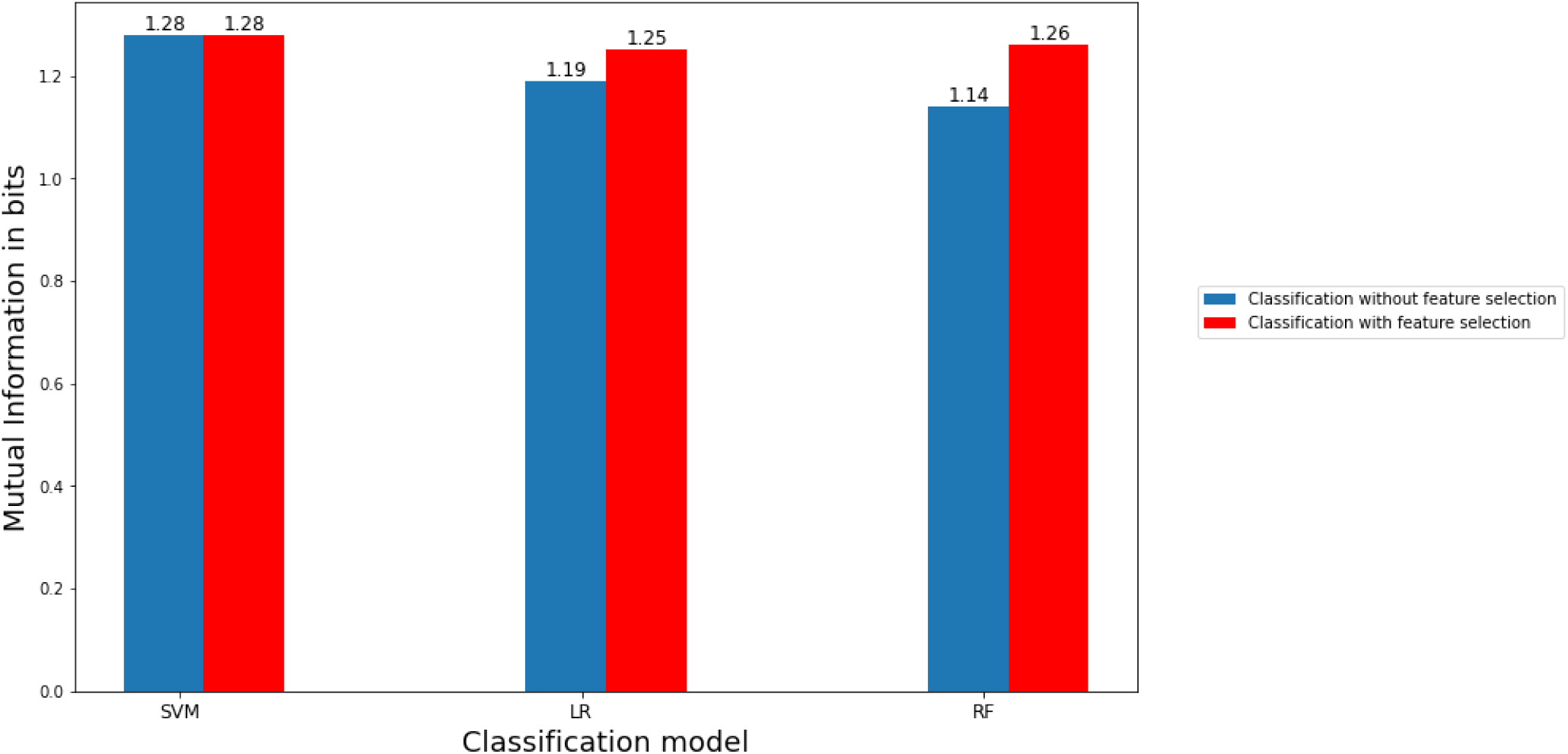
Mutual information with and without feature selection. The bar graphs display a comparison between the values of mutual information in bits between predicted and actual labels for the three models and for classification with feature selection and without feature selection as indicated

### Classification with randomly selecting 336 features

In order to test whether the GA based feature selection algorithm is able to select the features appropriately, We randomly chose 336 features from our dataset and evaluated the prediction accuracy using the SVM, LR, and RF models. We achieved accuracy of 0.83, 0.83, and 0.83 for the models. Table 6 shows the f1 scores of the different classes for the three models.

**Table 6:** F1 scores of different models of the different classes with randomly selecting 336 features

| F1 scores |  | SVM | LR | RF |
| --- | --- | --- | --- | --- |
| Malaria Life Cycle Stage |  |  |  |  |
| early_troph |  | 0.69 | 0.70 | 0.68 |
| late_troph |  | 0.86 | 0.85 | 0.87 |
| ring |  | 0.89 | 0.89 | 0.89 |
| schizont |  | 0.77 | 0.72 | 0.75 |

The accuracy and the f1 scores of this experiment were lower compared to the classification results with feature selection using the GA pipeline. This results demonstrate the legitimacy of the feature selection method.

### Construction and analysis of protein-protein interaction network

Understanding protein-protein interactions (PPIs) is critical for cell physiology in normal and pathological states because they are required for practically every process in a cell. ^15^ Protein-protein interaction networks (PPIN) are graphs of the physical interactions between proteins in a cell. Protein-protein interaction happens in specified binding areas and serves a specific function. The feature selection method provided us with 336 proteins in Plasmodium falciparum. We used the Search Tool for the Retrieval of Interacting Genes/Proteins database (STRING 11.0b)^16^ to construct the PPI network associated with these proteins. STRING can then construct a PPI network containing all of these proteins and their connections. Their interactions were generated with high confidence from high-throughput lab experiments and prior information in curated databases (sources: experiments, databases; Scores 0.90). The network construction shows a set of highly connected modules (Figure 7).

**Figure 7:**
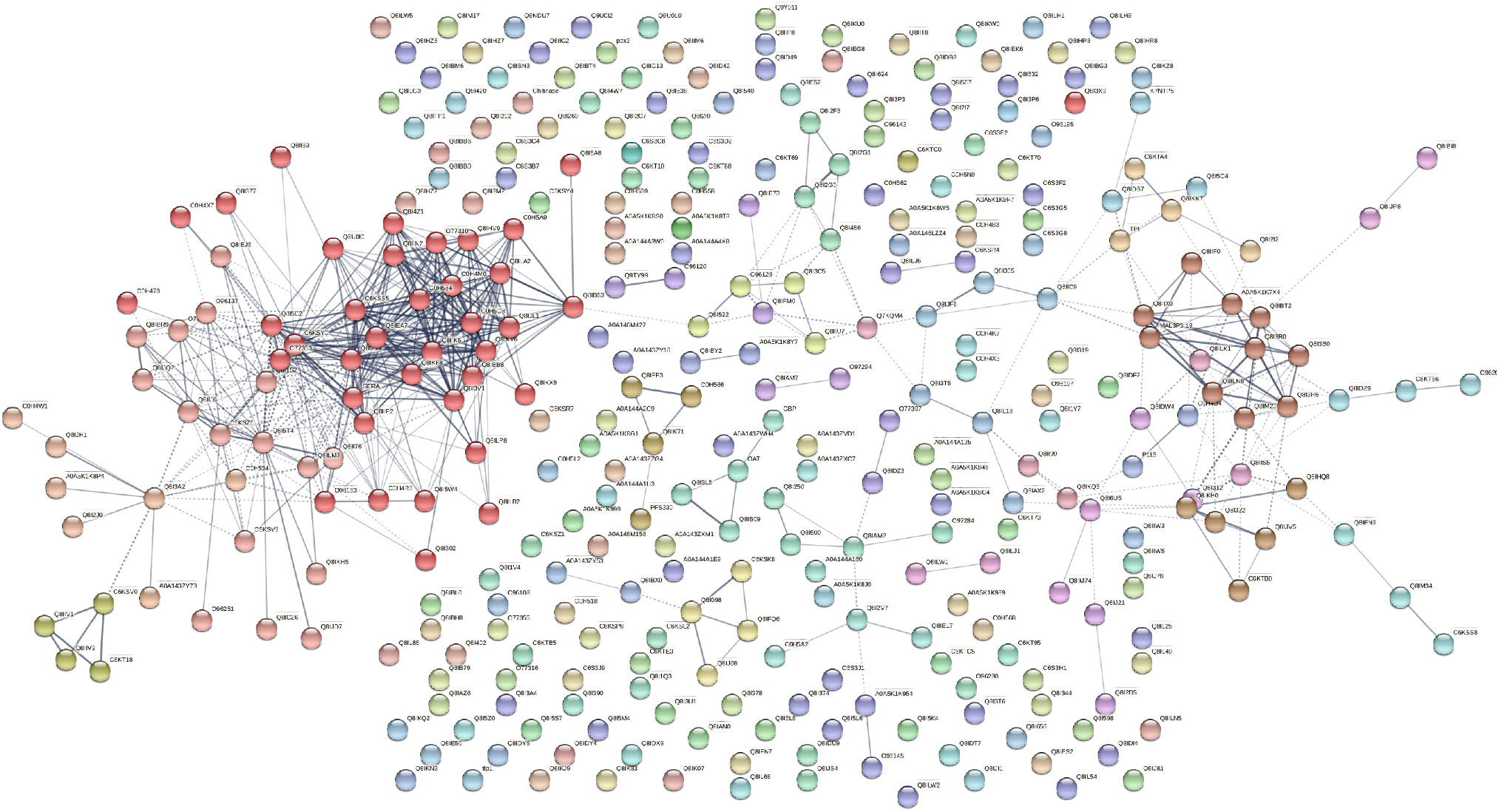
Protein-protein interaction network exhibits different clusters. The graph shows the protein protein interaction network of the 336 proteins selected by the feature selection method. The different colors indicate different identified clusters.

### The topological analysis of the PPI network

Various topological measures are generally used to evaluate the both global and node characteristics in the PPI networks, including degree (k), between centrality (BC), eccentricity, closeness centrality (CC), eigenvector centrality (EC), and clustering coefficient. ^17^ Here, highest degree nodes are identified using degree distribution. Additionally, We have used Markov Clustering Algorithm (MCL) to find clusters in the network. Among these clusters, we identified clusters which also contains the node with highest degree and high betweennss centrality.

This PPIN is composed of of 334 nodes with number of edges: 571, average node degree: 3.42, avg. local clustering coefficient: 0.304, expected number of edges: 519 PPI enrichment p-value: 0.013. We can see that proteins in red cluster have highest degree and high betweenness centrality. So, we can consider red cluster as disease module. We have analysed other topological properties like degree, BC, eccentricity, CC, EC, clustering coefficient, etc of this Red cluster using Gephi. ^18^

The proteins in Table 7 from red cluster have high degree and betweenness centrality (BC). In this cluster, we have, number of nodes: 37, number of edges: 272, average node degree: 14.7 avg. local clustering coefficient: 0.79, expected number of edges: 128 PPI enrichment p-value: ¡ 1.0e-16. We can see that this cluster has lesser nodes with high interaction and high clustering coefficient. So, this is a small world network. We can see from the above table that C6KSS5, Q8I2V4, C6KSY0 and Q8IKV6 has highest degree with high BC. We considered these four proteins as the hubs or bottlenecks as this nodes has high degree (k) and BC. We have chosen 7 more proteins that has high degree and BC to consider them backbone of the PPIN. These proteins are C6KSS5, Q8I2V4, C6KSY0, Q8IKV6, Q8IIK5, Q8IEA7, Q8I3V1, C0H5C8, C0H584. These 9 proteins are highly connected in PPIN and has control over the network. Next we have checked for the functionality of all these proteins.

**Table 7:**
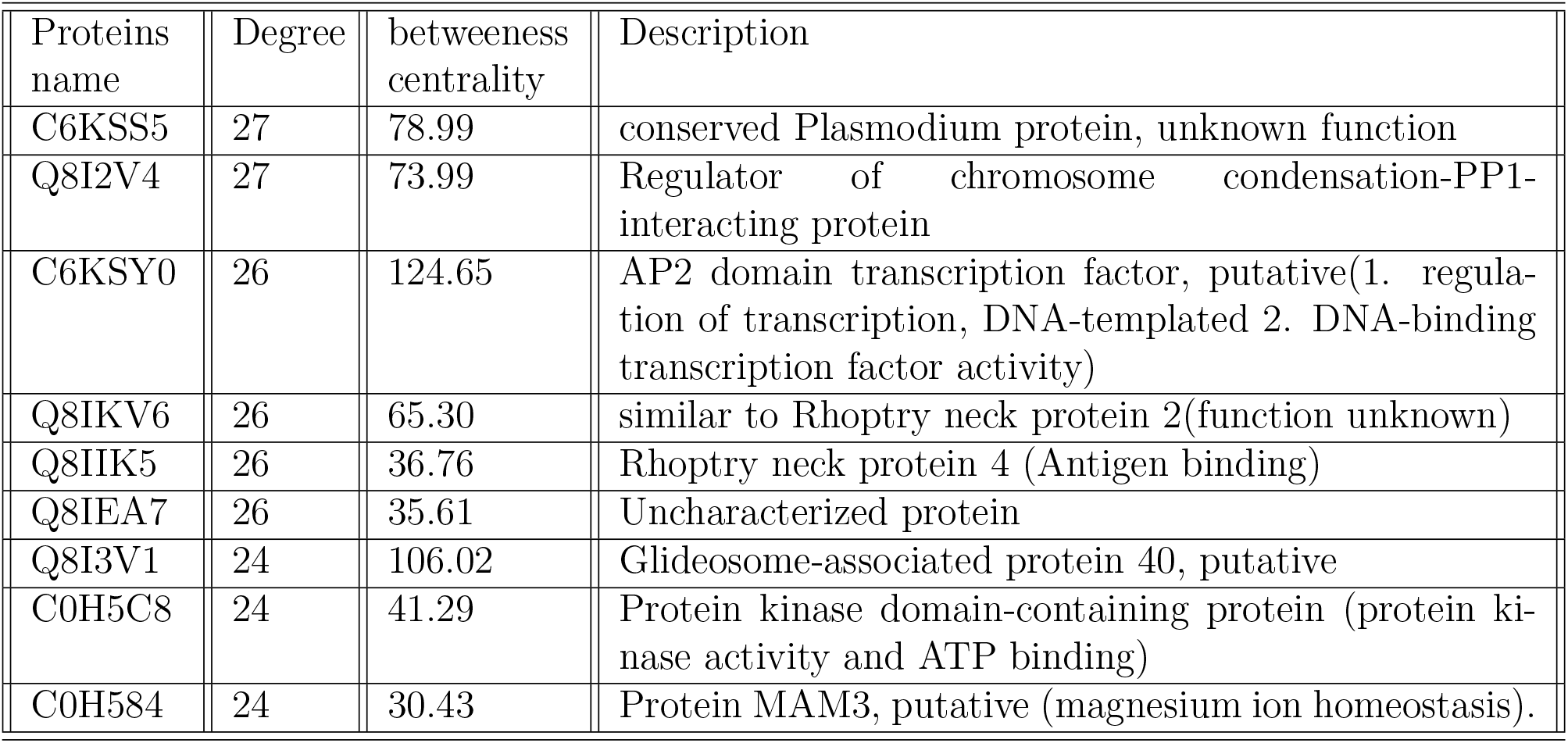
Topological analysis of the PPI network of the selected proteins

### Expression profile of the selected features

The analysis above provides us a set of proteins which are associated with the progression of the malaria pathogen through different stages of the life cycle. Thus, the expression pattern of these proteins would elicit the identity of the stages. In order to investigate the overall expression pattern of the genes across the different stages, we extracted the selected 336 features from the dataset. For each feature, we find the average RNA-seq read counts for all the four classes(early_troph, late_troph, schizont and ring). The average values are then transformed into log scale. We observed that genes fall into different clusters according to the expression patterns (Figure 8) and also the expression patterns vary among the stages. For instance, the genes at the bottom have a very low expression in ring phase. Similarly, genes at the top cluster are displaying low expression for all stages. These expression patterns may be harnessed to look for specific markers for different stages. Additionally, we visualised the clustering behaviour of cells after feature selection by GA, using the 336 features via the Seurat package. As done previously,the normalized counts were subjected to linear and non-linear dimensionality reduction using PCA and UMAP respectively. Figure 8b shows clear clusters of all the four blood cycle stages - ring, early_troph, late_troph and schizont, which supports that the selected features can serve as markers for the respective stages.

**Figure 8:**
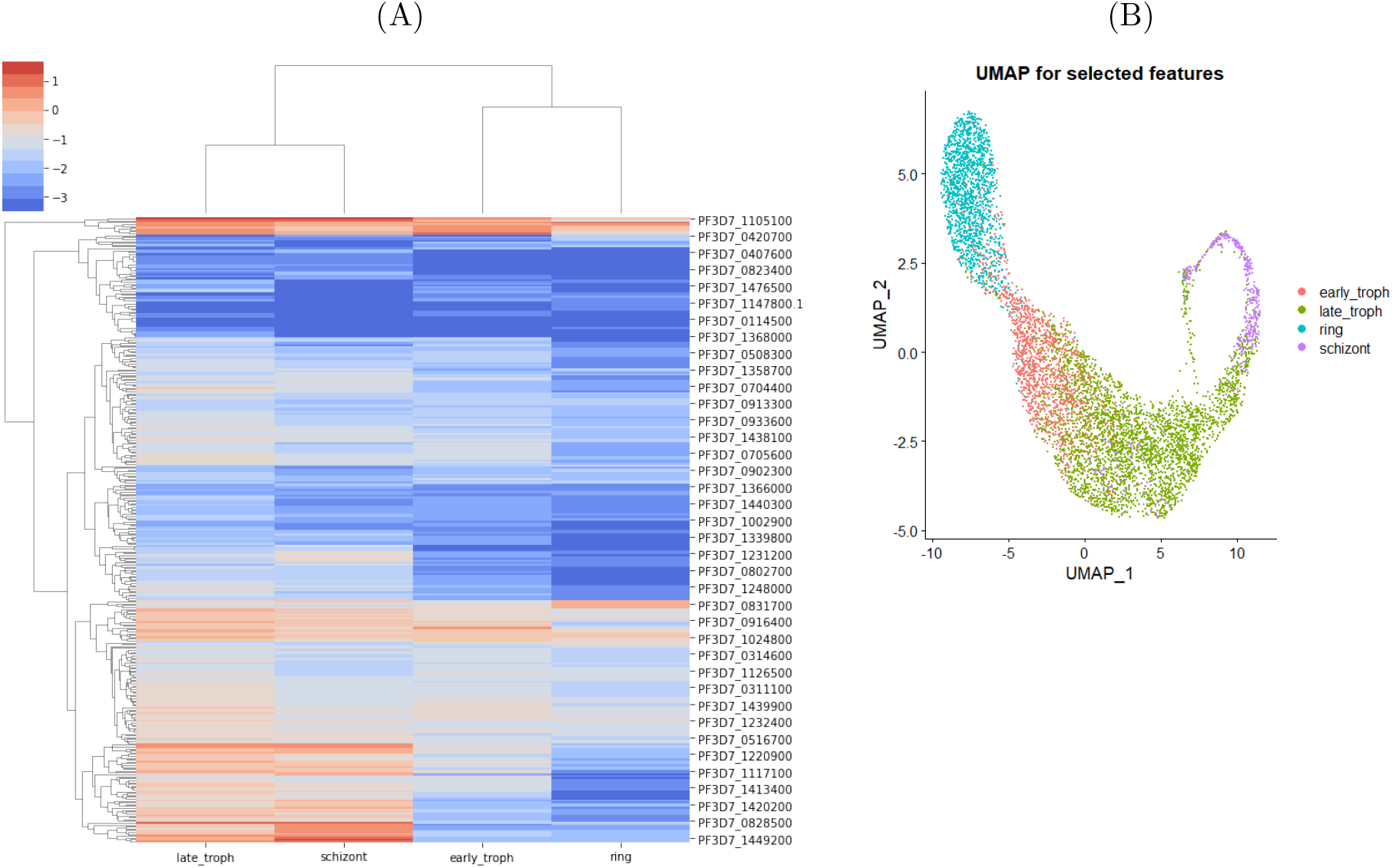
The expression profiles are distinct among the stages. (A) Expression Profile of the selected genes across the different stages. The heatmap shows the average RNA-count of the selected 336 genes across the different stages as indicated. A hierarchical clustering is performed on the expression levels in order group genes with similar expression patterns indicated by the dendogram. (B) UMAP of scRNA-seq counts of the selected features.Non-linear dimensional reduction of expression values of the selected 336 features.The cell clusters are colored based on the blood cycle stages of *P*.*falciparum*.

## Discussion and outlook

The present study has proposed a two-stage model for feature selection and classification which has been shown to improve the classification of the different stages of the Malaria Life Cycle. This was achieved by removing the irrelevant features from the total data set considered for analysis. The study’s main finding is that using a feature selection procedure before applying a classification algorithm results in more accurate predictions. The use of GA as a feature selection process significantly reduced the number of features included in the dataset.

Various machine learning (ML) approaches have been proposed to yield more accurate results. Karthik and Sudha, ^19^ reviewed ML methods for classifying gene expression model or computational analytical structure for complicated diseases, by identifying several differentially expressed gene techniques. Authors in ^20^ used the Convolutional Neural Networks (CNNs) based deep learning models for attribute extraction and categorization. For achieving higher categorization accuracy, they selected certain dominating features including size, colour, shape and cell count from the images. Similarly, a more effective two stage approach based on CNNs on a larger dataset was also proposed by.^21^ It remains an especially challenging task to distinguish the multiple growth stages of parasites. Seng et al.^22^ developed a deep learning approach for the recognition of multi-stage malaria parasites in blood smeared images using a novel deep transfer graph convolutional network (DTGCN). They reported higher accuracy and effectiveness compared to a wide range of state-of-the-art approaches.

Numerous ML approaches have been proposed in the literature to enhance gene expression data classification such as clustering, classification, dimensional reduction, among others ^23^. Training of ML models using initial high-dimensional features performs unsatisfactorily in practice and may result in network over-fitting and increased redundant information. This problem was addressed using random forests classifier in.^24,25^ Hossain et al. ^26^ designed an effective variational quantum circuit (VQC-based) approach to recognize the existence of malaria from RBC images through the classification of optimized feature set extracted from them. Murad et al. ^27^ used algorithms based on multifilter and hybrid approaches to feature selection leading to accuracy in excess of 90%. Mei et al.^28^ suggested a dimensionality reduction method for classifying tumour gene expression data. Arowolo et al. ^29^ and Li et al. ^30^ proposed a dimensionality reduction approach for classifying gene expression.

To overcome the dimensionality problem, Rokach et al. devised a genetic algorithm-based feature selection method. They evaluated the fitness function of several, obvious tree classifiers using a new encoding approach.^31^ Zhang et al. proposed a classifier ensemble with feature selection based on GA. The authors of this work created a new hybrid method that combines a multi-objective genetic algorithm with an ensemble of classifiers. The GA-ensemble approach was tested on a variety of datasets and its performance was compared using a variety of classifiers. ^32^ Cheng-Lung Huang ^33^ suggested a feature selection method based on GA and SVM optimization. The ultimate goal was to improve the SVM classification accuracy while optimising the feature subset and parameters. Chaung et al. ^34^ employed a hybrid technique that began with a genetic algorithm with a dynamic variable to pick a sample of genes, which were then ranked using chi square analysis, and the level of accuracy of the selection was assessed using SVM. Shutao et al^35^ used Particle Swarm optimisation and GA in order to perform highly accurate classification. The authors in ^36^ achieved a classification accuracy of 90.32 % using GA for feature selection and a SVM Classifier.

For further research the hybrid methods for feature selection, impact of parameter fine tuning on various algorithms’level and use of other methods including Ensemble Learning may be attempted. Next, we have constructed protein-protein interaction network between our proteins (which we get after using feature selection algorithm) and we have done network analysis on this protein-protein interaction network using STRING 11.0b and Gephi software. Various topological measures are estimated to evaluate the node characteristics in the PPI networks, including degree (k), between centrality (BC), eccentricity, closeness centrality (CC), eigenvector centrality (EC), and clustering coefficient. ^17^ We found degree and betweeness centrality of each protein though this calculation. Proteins having high degree and betweeness centrality tend to assert more control over the network function. These proteins can be considered as drug targets for future studies.

## Material and Methods

### System Design

The proposed classification technique comprises of two stages, namely:

1. Dimensionality Reduction with Feature Selection
2. Classification

### Dimensionality Reduction with Feature Selection

The dimensionality of the gene expression dataset is high. The dataset has redundant features which act as noise while training a model. This results in poor classification performance and high computational time. Dimensionality reduction is a technique that removes redundant features that hinder performance. We have used the feature selection ^37^ dimensionality reduction technique.

Let X be the initial m dimensional set of features, defined by the equation X = x(i)— i = 1,2,…m where x(i) are the defined features and m are the genes. The process of feature selection generates Y(i)—i = 1,2,…,p where Y(i) represents the new subset of features and p is now the number of features in the subset with p ¡= m. There are three types of feature selection methods - Filter, Wrapper, and Embedded approaches ^38,39^.

Figure 9 shows the the proposed computational model framework. The feature selection stage uses GA for dimensionality reduction.Genetic Algorithm (GA) is a metaheuristic, evolutionary, stochastic optimization algorithm inspired by the process of natural selection. GAs are commonly used to generate optimum solutions to problems employing three biologically inspired operators selection, crossover and mutation.

**Figure 9:**
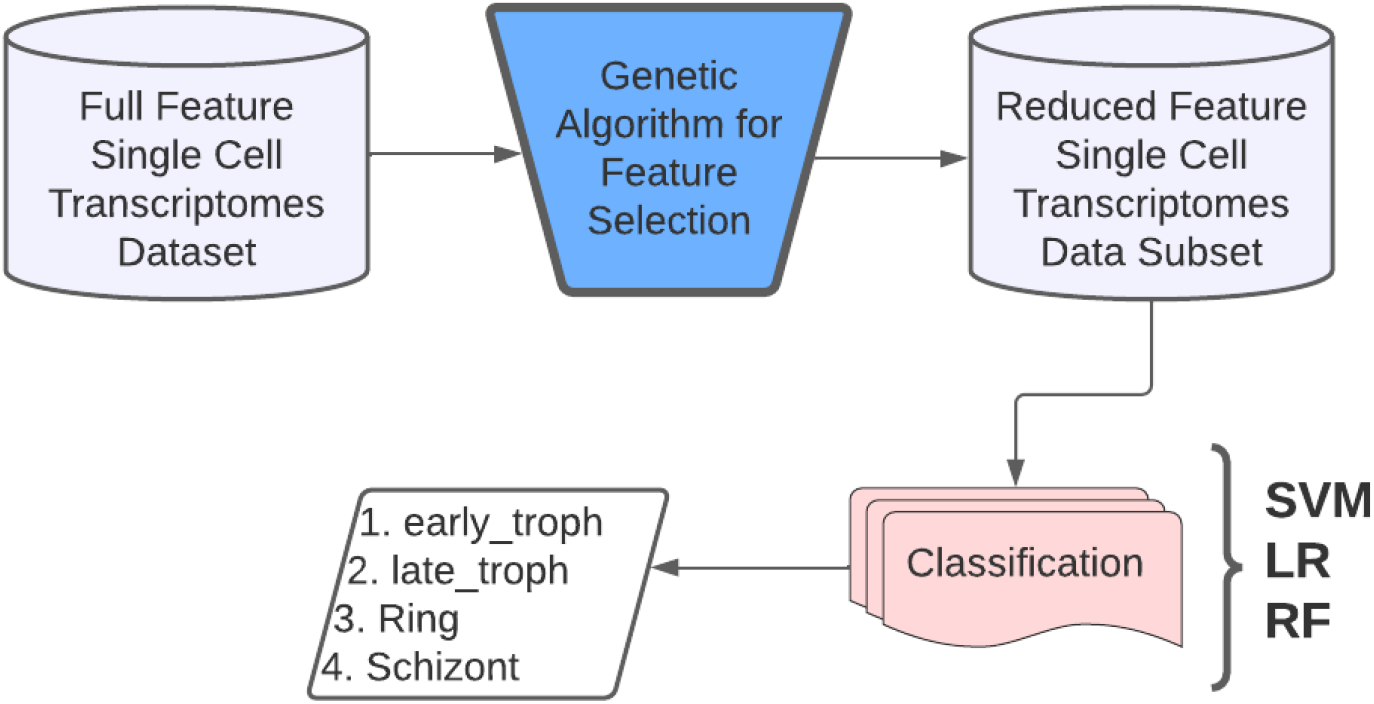
Proposed computational model framework ruled Comment/* */

### Classification

Classification is the process of identification of which of a set of categories or sub-populations an observation belongs to. Usually, the individual observations are grouped into a set of quantifiable properties called features. These features may either be categorical or ordinal or integer or real-valued. In the field of machine learning, the observations are called instances, the variables termed as features are grouped to form a feature vector, and the to be predicted categories are called classes. Figure 10 shows the pseudo code of the entire pipeline. We have used three classification algorithms in our research viz. Support Vector Machines, Logistic Regression and Random Forest.

**Figure 10:**
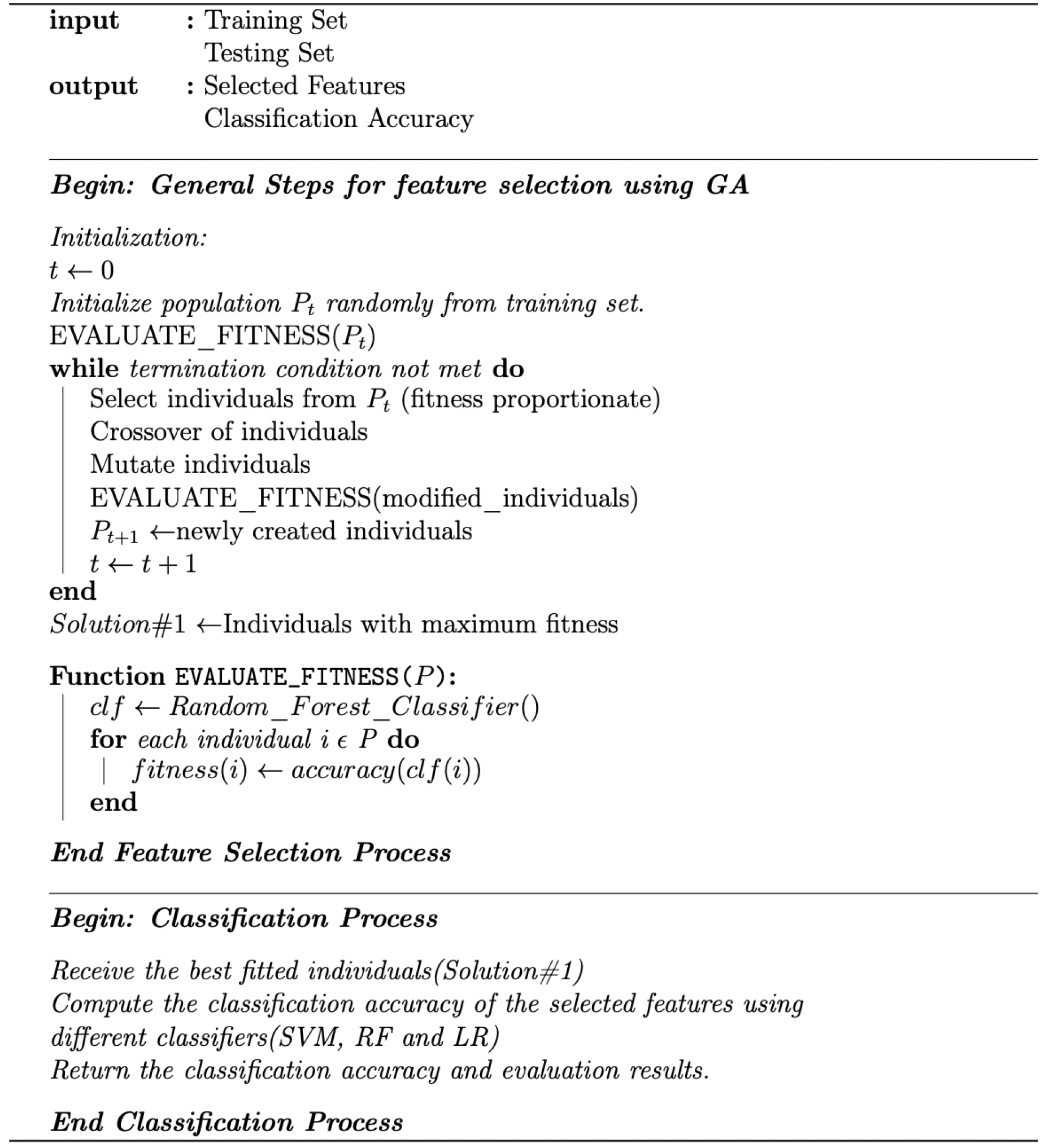
Psuedocode of the proposed method

Support vector machines (SVM) are one of the most popular, robust, non-probabilistic, binary, linear and non-linear classifier, supervised learning algorithms that analyze data for classification or regression analysis in machine learning. Based on a set of categorized training data, an SVM training model assigns new unseen examples to either of the trained categories, creating the decision boundaries (or hyperplanes) that can segregate n-dimensional space into classes.

Logistic regression (LR) is another powerful supervised ML algorithm used for binary classification problems that can be generalized to multiclass classification. A logistic function is used to model the probability of a discrete outcome based on an input variable. LR is an extensively employed algorithm for classification problems in industry owing to its high simplicity and efficiency particularly for linearly separable classes.

Random forests (RF) are an ensemble learning classification method and work by constructing a multitude of decision trees at training time. For classification jobs, the RF output is the class selected by most trees. Ensemble learning combines many classifiers to provide solutions to complex problems. In machine learning RFs also assist in reducing the training set overfitting by decision trees and also increases precision.

## Experimentation

The following sections will introduce each of the followed steps in detail. A summary of the implementation of the entire pipeline is depicted in Figure 11.

**Figure 11:**
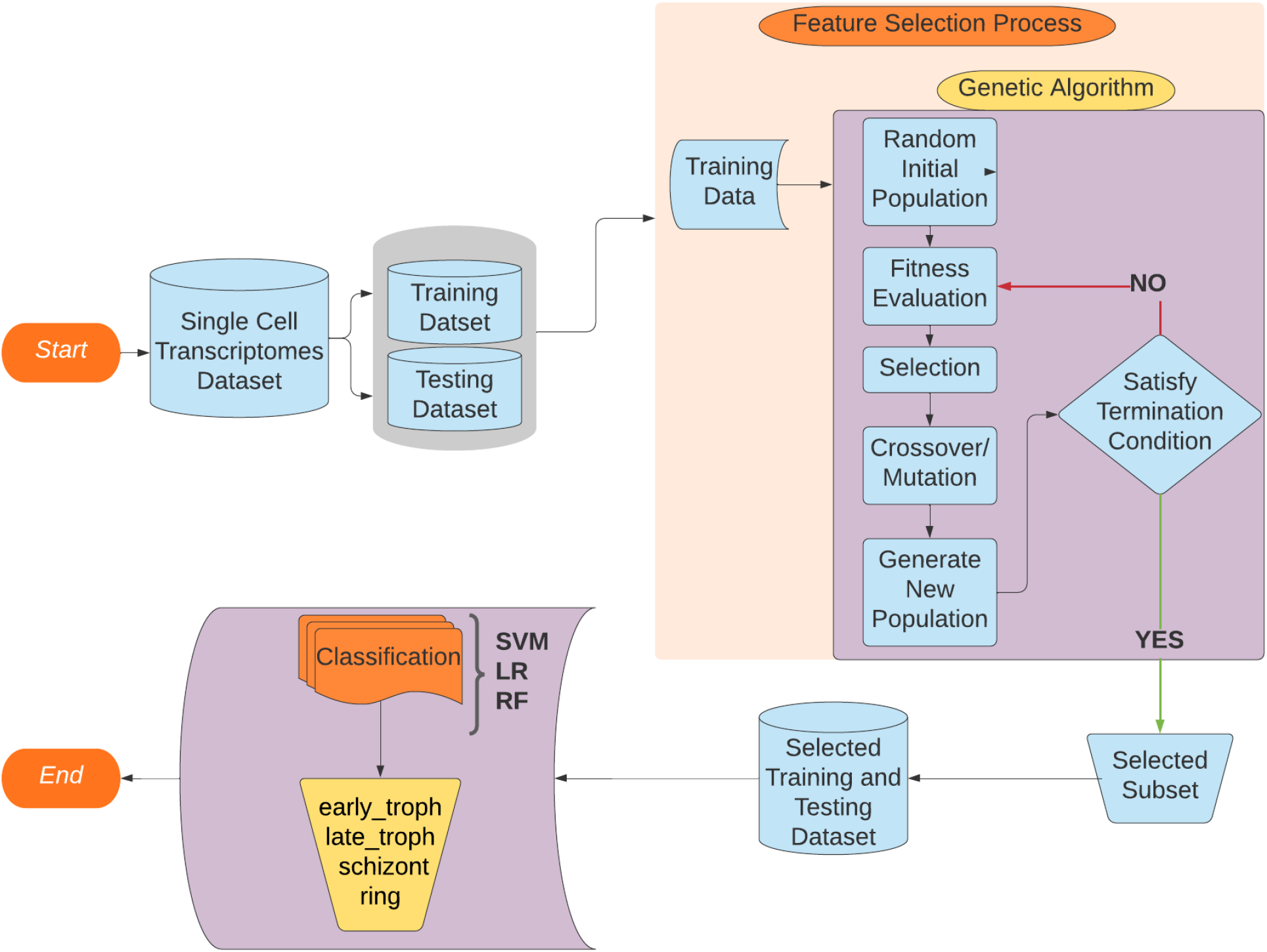
Flowchart of entire pipeline

### Data Preparation

In order to create an independent test set and improve the classification validity and accuracy, the input data was divided into the training and testing sets in a ratio of 80% and 20% respectively. The training set was created to validate the feature selection while the test set served a similar validation role in the classification process. The training set is then processed through the GA pipeline.

### Genetic Algorithm

Genetic Algorithm (GA) is a stochastic evolutionary optimization technique. It starts with an initial randomized set of population of features (500 here) and then creates another population using subsets of the available features whose individuals are evaluated using the Random Forest predictive model for the target task. The tournament selection technique is used to pick the higher fitness subsets to be carried forward into the next generation for applying the cross-over (updating the winning feature sets with features from the other winners) and mutation (probabilistically introducing or removing some features) genetic operators. The individuals of this subset are stored in the Hall of Fame which is continuously sorted so as to have the first element with maximum fitness value so far. This process is iterated to yield the optimum features for the set termination criteria (maximum generations = 100, if no change in the best-fitted individual for 20 generations). Few other GA modelling parameters used in our research are - uniform crossover probability of 0.5, flip-bit-mutation probability of 0.2.

### Classification Process

After the GA has selected the optimal features, these features are then subjected to different classification algorithms (SVM, Random Forest, Logistic Regression) to measure the classification accuracy of the selected feature set. This yields us the classification accuracy of the four different classes viz **early_troph, late_troph, schizont and ring**.

### Mutual Information

Mutual Information (MI) is a measure of how much one variable’s uncertainty is reduced when the other variable’s value is known. It is given by the formula: ^40^

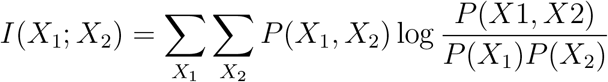

where *P*(*X*_1_, *X*_2_) is the joint distribution of the two variables. *P*(*X*_1_) and *P*(*X*_2_) is the marginal distribution of the two variables. It’s a dimensionless quantity that’s measured in bits. Each element of confusion matrix represents the conditional probability of predicting a class y’ given the true category y - *p*(*y*^*′*^|*y*). The joint probability of *P*(*y, y*^*′*^) is equal to the multiplication of probability of true label *P*(*y*) and conditional probability *P*(*y*^*′*^|*y*). *P*(*y*^*′*^) is given by sum of joint probability over true label y. We have used this to find the *I*(*y*; *y*^*′*^).

## Data and Software Availability

The data are freely accessible as a processed dataset through a user-friendly web interface (www.sanger.ac.uk/science/tools/mca/mca/).^13^ Our dataset has 5066 rows and 6737 columns. Each row corresponds to single cell and each column corresponds to a gene. We have 5066 feature in our dataset. We have four different malaria life cycle stages (early_troph, late_troph, ring, schizont).

**[Ada](https://www.iiit.ac.in/hpc/index.html)**, the High Performance Computing Data Center of International Institute of Information Technology Hyderabad, India was utilized for the computation. It consists of 92 nodes, each equipped with dual Intel Xeon E5-2640 v4 processor, 128 GB RAM, two scratch disks (2 TB SATA and 960 GB SSD SATA) and four Nvidia GTX 1080 Ti / RTX 2080 Ti GPUs. The cluster has a total of 1472512 GPU cores, 3680 CPU cores and 11776 GB RAM. For our experiment we have used 40 cores with maximum memory per CPU as 2 GB on a Linux Ubuntu operating system. The proposed model is implemented using Python with the genetic selection library for the Genetic Algorithm implementation and the sklearn library for the classification algorithms. The relevant data and python scripts can be found in this github **[code](https://github.com/swarnimshukla/Machine-learning-approaches-for-classification-of-Plasmodium-falciparum-life-cycle)** link. We used the R-based Seurat (v4.1.0) package developed by Satija lab^14^ for visualisation and dimensionality reduction of single cell RNA-seq data. This was implemented in R (v4.1.3), run on RStudio environment (v1.3.1093). We followed the standard pre-processing workflow, normalisation, linear and non-linear dimensionality reduction recommended by Seurat developers with default parameters, unless otherwise mentioned in the results section. The feature selection method provided us with 336 proteins in Plasmodium falciparum. We have used the Search Tool for the Retrieval of Interacting Genes/Proteins database (STRING 11.0b)^16^ to construct the PPI network associated with these proteins. STRING software https://string-db.org/ can then construct a PPI network containing all of these proteins and their connections. Their interactions were generated with high confidence from high-throughput lab experiments and prior information in curated databases (sources: experiments, databases; Scores 0.90). Various topological measures are generally used to evaluate the both global and node characteristics in the PPI networks, including degree (k), between centrality (BC), eccentricity, closeness centrality (CC), eigenvector centrality (EC), and clustering coefficient. ^17^ Here, highest degree nodes are identified using degree distribution. Additionally, We have used Markov Clustering Algorithm (MCL) (using STRING) to find clusters in the network. Among these clusters, we identified red cluster which contain the node with highest degree and high betweennss centrality. We have analysed topological properties like degree, BC, eccentricity, CC, EC, clustering coefficient, etc of the Red cluster using Gephi^18^ software.

## Acknowledgement

Authors thank the Department of Biotechnology (No. BT/RLF/Re-entry/32/2017), Government of India for funding.

